# Functional annotation hypothetical proteins: a world to be explored in drug development in Trypanosomatids

**DOI:** 10.1101/2023.12.07.570673

**Authors:** Raissa Santos de Lima, Ana Carolina Silva Bulla, Manuela Leal da Silva

**Affiliations:** Programa de Pós-Graduação em Biologia Computacional e Sistemas, Instituto Oswaldo Cruz, Manguinhos, Rio de Janeiro, RJ, Brazil; Instituto de Biodiversidade e Sustentabilidade NUPEM, Universidade Federal do Rio de Janeiro, Macaé, RJ, Brazil

## Abstract

Hypothetical proteins can provide an alternative pathway for finding potential targets in the development of new drugs due to the fact that many Neglected tropical diseases are caused by Trypanosomatids (Chagas Disease, Leishmaniasis, and Human African Trypanosomiasis). In this work, we focus on applying functional prediction methods based on both sequence and structure to analyze the hypothetical proteins of the pathogenic agents that cause these diseases: *T. cruzi* (Tcr), *T. brucei brucei* (Tbr), *T. brucei gambiense* (Tbg), *L. infantum* (Lif), *L. donovani* (Ldo), and *L. braziliensis* (Lbz). By consulting databases and servers, we have predicted functional domains for twenty-six proteins in Tcr, thirteen in Tbr, fifteen in Tbg, ten in Lif, and one in both Ldo and Lbz. With the goal of developing multi-target therapies, we grouped the domains according to how they are shared among the organisms and investigated those that are shared among more species. By examining the existing literature using specific search strategies, we described what has already been reported for these domains and also analyzed protein structures and sequences, describing mutations among the species and potential drug sites. The published works have unveiled that some of these domains are non-essential for trypanosomatids, like the TRX domain, while others demand further investigation due to a lack of information about metabolic processes (UFC1, Ufm1, ACBP, AAA 18, and Fe-S). Although, we have identified three noteworthy domains that hold promise as targets: TPR, which plays a crucial role in the ciliogenesis process; Nuc deoxyrib tr, essential in purine recycling and recovery mechanisms; and MIX, important for protein targeting and the assembly of complexes such as COX. These three domains are promising targets for drug development due to their conservation, their potential to affect multiple species and their exclusivity.

## Introduction

Trypanosoma parasite species can infect several animals causing neglected diseases such as Chagas disease (CD) (*Trypanosoma cruzi*), Human African Trypanosomiasis (HAT) (*Trypanosoma brucei*) and Leishmaniasis (genus Leishmania) [1,2]. From an epidemiological standpoint, these diseases’ incidence rates are alarmingly high, with millions of cases reported annually [3–5]. Furthermore, HAT was close to eradication in the 60s but the disease has since experienced a resurgence in reason of political problems, and a decline in control programs [5,6]. Besides that, many of these diseases have a wide geographical distribution due to infected individuals immigrating to non-endemic countries, and leishmaniasis in particular has three endemic foci: India, East Africa, and Brazil [3,7,8].

Aside from the challenge of dissemination, the treatment has several problems, mainly regarding efficacy and therapeutic adherence due to the high toxicity of the drugs used [9–11]. Furthermore, there are already cases of resistance that demonstrate the promiscuity not only of drugs but also of current targets for diseases caused by trypanosomatids [12,13]. As a result, there is a pressing need to take the initiative to develop new drugs, and one promising approach is the drug repositioning strategy. This process is highly cost-effective since it reduces drug development time, optimizing pre-clinical trials by utilizing drugs with known possible adverse effects [14,15]. Thus, efforts are required to identify new molecular targets for new drugs and their implementation as effective treatments.

Drugs can have different molecular targets, one of the most common being proteins, and with the increasing popularity of sequencing techniques, there has been a rise in the use of bioinformatics tools, including those for protein analysis [16]. Despite the availability of predicted proteins in genomes, there is a limited amount of functional details data for these molecules. The reason for that is the sheer volume of data that experimental functional characterization produces, which is difficult to manage. So the functional prediction approach comes in as a solution, and it has been constantly improved [17].

There are two main approaches to protein function prediction and the first one is the SEquence-Based Functional Prediction (SEBFP), which assumes that the sequence contains a conserved “functional signature” that can be identified even in a small part of the sequence [18]. Consequently, if two proteins have this conserved “signature” it is possible to infer the protein function [19]. The second approach is Structure-Based Functional Prediction (STBFP) which has two premises, the first one is that all the essential information about a protein’s structure is contained in its amino acid sequence [20]. The second premise is that proteins have a specific native folding state or a set of native states, and this structure is closely related to the catalytic protein function [21].

The primary aim of this study is to employ a hybrid approach, merging SEBFP and STBFP strategies, to predict the functional domains of hypothetical proteins (HP) as depicted in Fig 1. Our goal is to achieve more robust results compared to conventional linear methodologies and to identify new targets for drug development that are applicable across multiple neglected diseases caused by these parasites.

**Fig 1.**
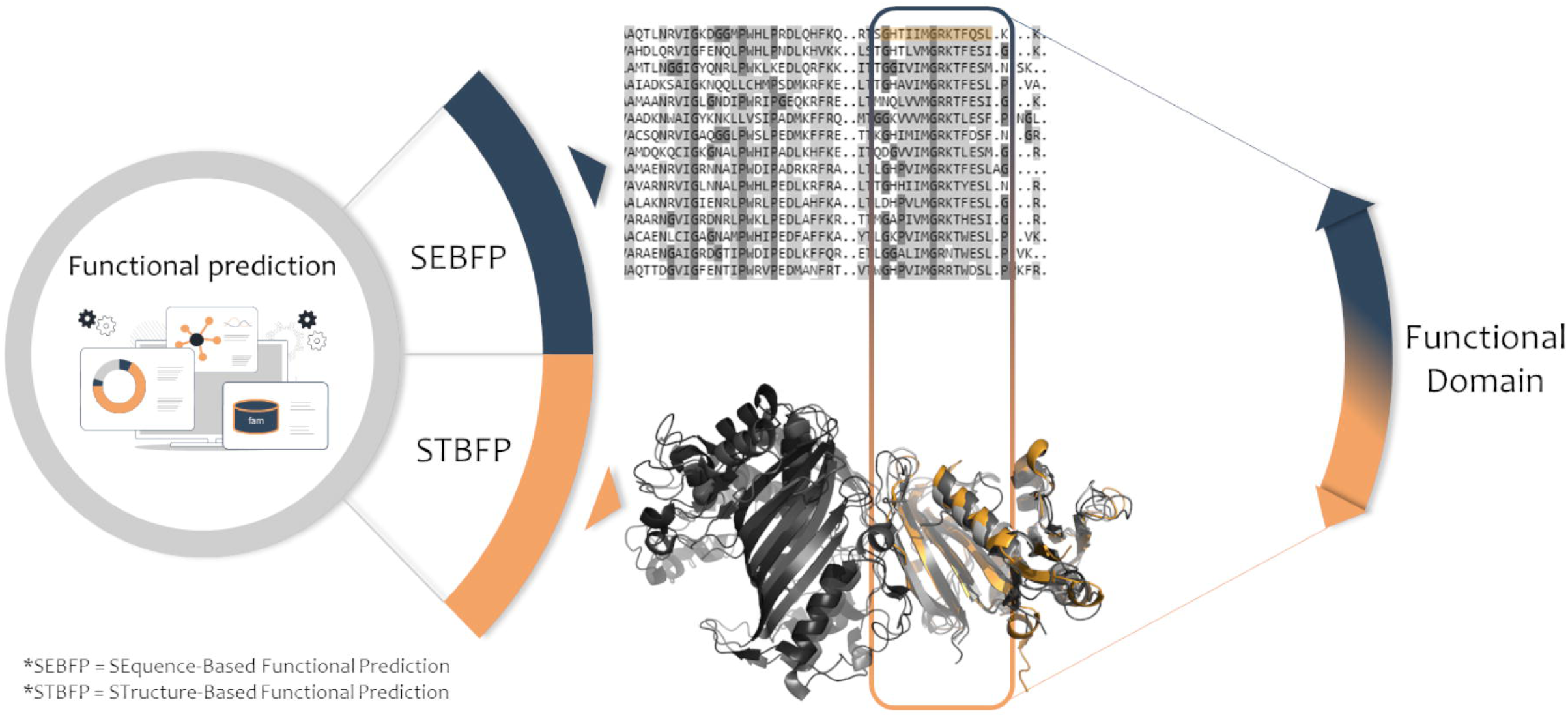
Scheme of the functional prediction methodology. Through the utilization of a diverse range of computational tools and databases, we conducted an extensive analysis of both sequence and structural data to identify functional domains within the hypothetical proteins (HP). As illustrated in the sequence and structure, an exemplary functional domain is highlighted in orange, showcasing its conservation across various protein data levels. For the 3D protein representation, the PDBids were used: 3UM8, 4TU5, 6WEP, and 3Q1H.

To demonstrate our approach, we applied SEBFP and STBFP using various databases, identifying a total of sixty-six protein domains for the organisms *T. cruzi* (Tcr), *T. brucei brucei* (Tbr), *T. brucei gambiense* (Tbg), *L. infantum* (Lif), *L. donovani* (Ldo), and *L. braziliensis* (Lbz). Notably, nine of these domains were shared by four species (Tcr, Tbr, Tbg, and Lif), which we have provided detailed descriptions for. With a focus on identifying promising multi-targets, we conducted sequence and structural analyses in our pursuit of potential druggable sites for the three domains (TPR, Nuc deoxyrib tr, and MIX) that hold promising functions.

## Materials and methods

### Proteomes selection

According to the disease, we selected the proteomes of the following species *Trypanosoma cruzi* for CD, *Trypanosoma brucei* for HAT and *Leishmania* spp for Leishmaniasis. To determine the strain among several deposited in the National Center for Biotechnology Information (NCBI) databases [22], we first selected the most widely reported strain based on literature data. Additionally, we considered the strains with the highest assembly level available in the NCBI. After the selection process, using a script that analyzes the identifiers, we separate the proteins annotated as HP from the others.

### Large-scale comparative modeling

The multiFASTA file containing only the HP sequences was submitted to the MHOLline web server for structural modeling [23]. MHOLline is a biological workflow capable of performing three-dimensional modeling for different proteins at the same time. The structure prediction process starts with searching for templates by aligning protein sequences with structures in the Protein Data Bank (PDB) using the Basic Local Alignment Search Tool (BLAST) [24]. Monomeric 3D models are generated with MODELLER [25] for only results with e-value ≤ 10e-5, identity ≥ 0.25, and Length Variation Index (LVI) ≤ 0. Subsequently, MHOLline classifies these models into five groups among Very High and Very Low using the module called Filters, which is based on the identity and LVI values. In this study, we selected those that were classified between Very High and Good (Very High: ID ≥ 75% and LVI ≤ 0.1%; High: ID ≥ 50% and <75% and LVI ≤ 0.1%; Good: ID ≥ 35% and < 50% and LVI ≤ 0.3%). Hence, only the protein sequences that met the evaluation criteria were deemed suitable for the functional prediction stage.

### Functional prediction and description

As previously mentioned, we employed a combined SEBFP and STBFP approach to predict the selected HP sequence’s functional domains. In total, nine databases and six web servers were utilized, as shown in Fig 2. To ensure the reliability of the functional prediction, we defined that the same domain must be predicted in at least eight of the SEBFP tools and two of the STBFP for functional inference. Similar strategies have been previously adopted by Shahbaaz, M., Hassan, M. I., & Ahmad, F. (2013) and Mazandu, G. K., & Mulder (2008).

**Fig 2.**
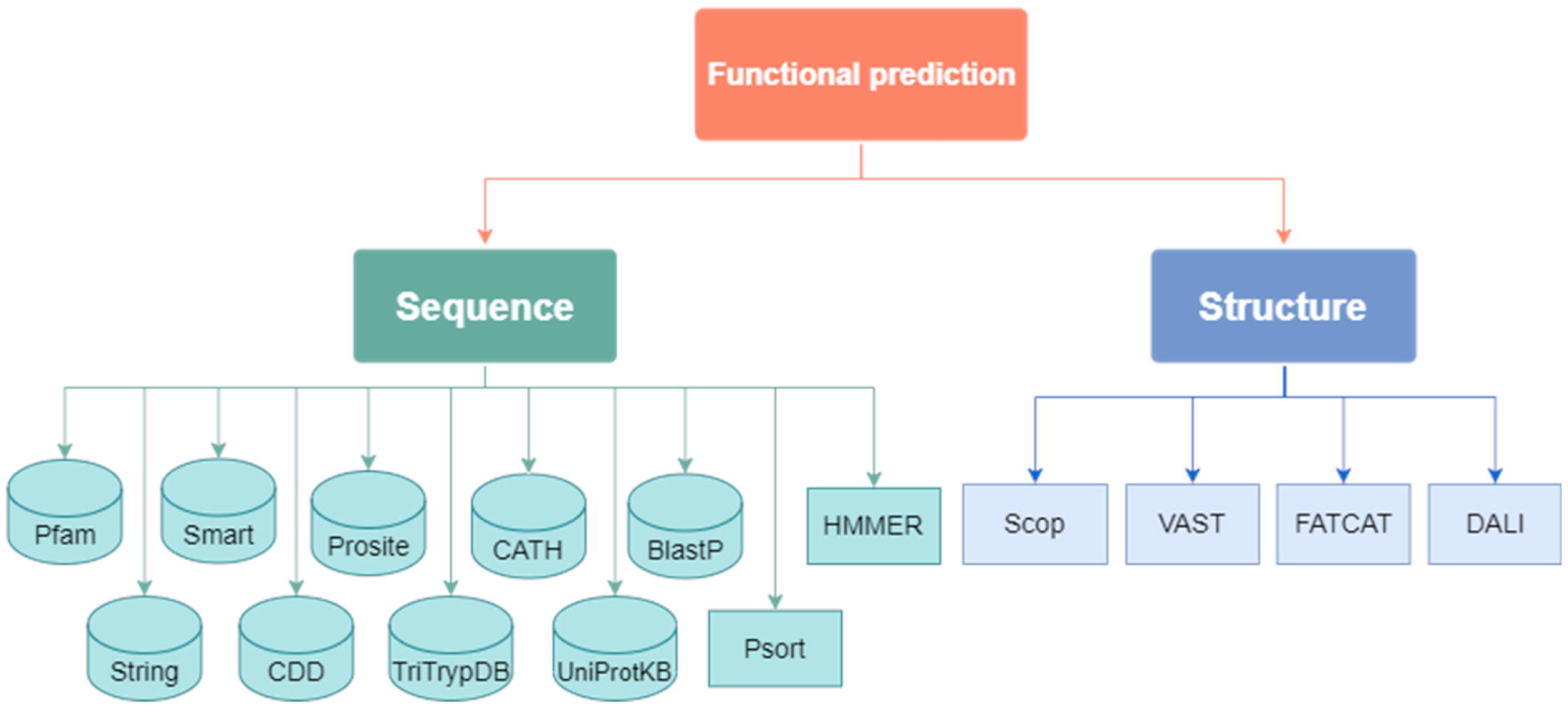
Workflow containing databases and servers used in SEBFP and STBFP. Databases are represented as cylinders and the web servers as rectangles.

Consulting the literature, the functional domains were detailed, particularly through the use of bibliographic databases such as Web of Science, Google Scholar, The Lens, and PubMed. The search strategy employed was using keywords such as the functional domain acronym, disease’s name, and the parasite species, for example, “TRX AND Chagas Disease AND Trypanosoma cruzi”. Additionally, to find targets that are crucial for the parasite’s metabolic pathways, we consulted the Kyoto Encyclopedia of Genes and Genom3es (KEGG) database [26] using the EC number or the functional domains acronym in the.

### Sequence analysis

Sequence analysis began using tools that contain details about domains and families, which included: Pfam [27], Simple Modular Architecture Research Tool (SMART) [28], PROSITE [29], Conserved Domains Database (CDD) [30], Search Tool for Retrieval of Interacting Genes/Proteins (STRING) [31] and the CATH [32] and HMMER [33] web servers. These tools were used with default settings and allow the identification of homologous sequences by aligning them with information in various databases. This type of approach, the homologs identification, or evolutionary relationships, is widely used for the structural and functional annotation of proteins [32–34].

The proper characterization of a protein involves determining not only its function but also its cellular localization. This information is crucial for understanding its role in cellular processes and for the development of targeted therapies. To determine the subcellular localization we used the Psort (Prediction of Protein Sorting Signals and Localization Sites in Amino Acid Sequences) web server [35], more specifically WoLF PSORT with default parameters. This tool predicts protein sorting signals and localization sites based on amino acid sequences. Lastly, we consulted UniProtKB, where is possible to search for similar sequences with predicted functions using the integrated BLAST tool and we consulted the curated Swiss-Prot sequence database with default parameters [36,37].

To further explore sequence similarity with other sequences with other already known sequences deposited in data banks, we used the Basic Local Alignment Search Tool (BLAST) tool with the Non-Redundant (NR) [24] database and the TriTrypDB database [38], both with the default search parameters. The TriTrypDB database contains sequence data for the Trypanosomatidae family species and sources its information from banks such as the DNA Data Bank of Japan (DDBJ) and the European Molecular Biology Laboratory (EMBL) [39].

Among species, the domain conservation, as well as mutations were analyzed using the UGENE alignment tool, specifically with the MUtiple Sequence Comparison by Log-Expectation (MUSCLE) algorithm [40]. MUSCLE realigns the sequences repeatedly, considering a guiding phylogenetic tree to generate the distance matrix [40–42]. Given that we were analyzing organisms from the same family, considering the phylogenetic relationships when assembling the matrix was crucial for the alignment quality [43].

Finally, in addition to the aforementioned analyses, we identified the signal peptide and transmembrane regions. First, for signal peptide detection, we used the SignalP web server, which can predict not only the signal peptide’s presence but also their cleavage sites [44]. For the transmembrane regions encounter, the DeepTMHMM web server was used, which scores the protein region probability being inside, in the transmembrane region, or outside the membrane [45]. We used both servers with the default search parameters.

### Structure analysis

STBFP analyses started by utilizing the structures generated by MHOLline to search the SCOP server. This server classifies the protein based on its structure and provides a structural relationship description among known proteins [46]. Besides that, we used the VAST server, which compares the spatial coordinates of an input 3D structure with other structures deposited in the Molecular Modeling Database (MMDB) of NCBI [47]. Furthermore, VAST can identify distant homologs that may not be recognized through a simple sequential comparison [48].

Another server utilized was the Flexible structure AlignmentT by Chaining Aligned fragment pairs allowing Twists (FATCAT), which searches for similar structures in databases such as PDB and SCOPe through a structure alignment algorithm [49]. Lastly, the Distance matrix ALIgnment server (Dali) was used also for comparison structures in the PDB database but it utilizes three different ways: the heuristic search (all structures in PDB), the exhaustive search (a PDB subset), and an AF-DB hierarchical lookup (AlphaFold subset) [50]. Both FATCAT, Dali, and VAST servers perform alignment based on calculating of Root Mean Square deviation (RMSD) using α carbons.

Regarding the RMSD, the Pymol (version 1.9) program [51] was utilized to calculate it for the models generated by MHOLline with the templates used. In addition, we examined the conservation of amino acid residues that compose the functional domain, as well as the presence of mutations among the species in the structures. To evaluate regions without clearly defined secondary structures, we referred to the AlphaFold structure database [52], which provides information on models generated by artificial intelligence [53], including details on species. Moreover, we utilized the Quick2D web server, a tool that includes nine predictors of secondary structures, even regions of disorder, to facilitate reaching a consensus on the structure in the region of interest [54].

### Modeling homo-oligomeric structures and validation and binding site identification

The final structures were remodeled considering the oligomeric state through AlphaFold2 on ColabFold [55] pipeline (num_relax: 5, template mode: none, msa_mode: MMSeq2 (uniRef[+[environmental), num_recycle: 3), relaxing the structure with an amber force field, considering the sequences in table S4. To verify the structural quality of the built models, the models were submitted to MolProbity [56] and ProSA – web [57]. To access the protein druggability, the structures were submitted to DogSiteScorer [58] server for prediction of potential pockets on the proteins (score between 0 and 1).

## Results and Discussion

### Selection and large-scale 3D modeling

Using the NCBI database, we conducted a search of organism’s proteomes, specifically: *Trypanosoma cruzi* (Tcr) for CD; *Trypanosoma brucei brucei* (Tbr), and *Trypanosoma brucei gambiensi* (Tbg) for HAT; *Leishmania infantum* (Lif), *Leishmania donovani* (Ldo) *Leishmania braziliensis* (Lbz) for Leishmaniasis. Lif and Ldo species were selected due to their association with the most severe clinical form of the disease, visceral leishmaniasis (VL), which is more prevalent in Brazil [3,7].

The organisms and their respective strains were selected considering literature data and the proteomes with the highest assembly level deposited in the NCBI database (Table 1). Once the strains were chosen, the hypothetical proteins (HP) were separated from the others and submitted to the MHOLline server. This resulted in the generation of 3D models for 2928 proteins, ouch of which 111 models qualified for the functional prediction stage due to their classification between Very High and Good (Very High: ID ≥ 75% and LVI ≤ 0.1%; High: ID ≥ 50% and <75% and LVI ≤ 0.1%; Good: ID ≥ 35% and <50% and LVI ≤ 0.3%). Lastly, we consulted sequential and structural databases to infer functional domains for these proteins.

**Table 1.** Correlation of species, numbers of proteins, and 3D models.

| Species | Strains | NCBI Code | TP | HP | Gen 3D | Models selected |
| --- | --- | --- | --- | --- | --- | --- |
| *T. cruzi* | CLBrener | GCA_000209065.1 | 19.608 | 11.171 | 1.421 | 40 |
| *T. brucei brucei* | TREU927 | GCA_000002445.1 | 8.758 | 5.711 | 658 | 20 |
| *T. brucei gambiensi* | DAL972 | GCA_000210295.1 | 9.822 | 6.611 | 661 | 23 |
| *L. infantum* | Jpcm5 | GCA_000002875.2 | 8.15 | 5.245 | 648 | 21 |
| *L. donovani* | LdCl | GCA_003719575.1 | 8.632 | 3.527 | 90 | 3 |
| *L. braziliensis* | MHOM | GCA_900537975.1 | 8.244 | 3.373 | 98 | 4 |

<sup>a</sup>TP - Total of proteins
<sup>b</sup>HP - Hypothetical protein
<sup>c</sup>Gen 3D - 3D models generated

### Annotation of HPs

The nomenclature used to describe the results was based on the user in the following databases: InterPro [59], Pfam [27], SMART [28], PROSITE [29], and CDD [60] (S1 Table). To predict the functional domains, we established the following criterion: the same function must be predicted in at least eight SEBFP tools and two STBFP tools. Using sequence and structure analysis tools, we inferred 26 functional domains for Tcr, 13 for Tbr, 16 for Tbg, 10 for, Lif, 1 for Ldo, and 1 for Lbz, totaling 67 proteins from the initial 111 (S2 Table).

By analyzing SEBFP and STBFP results together, it was observed that some of the annotated domains were repeated within the same species and also interspecies (Fig 3). Specifically, we found that nine predicted domains were shared among four species of trypanosomatids, namely Tcr, Tbr, Tbg, and Lif. Therefore, we narrowed our focus to these shared domains, as our goal is to identify multi-targets for trypanosomatids.

**Fig 3.**
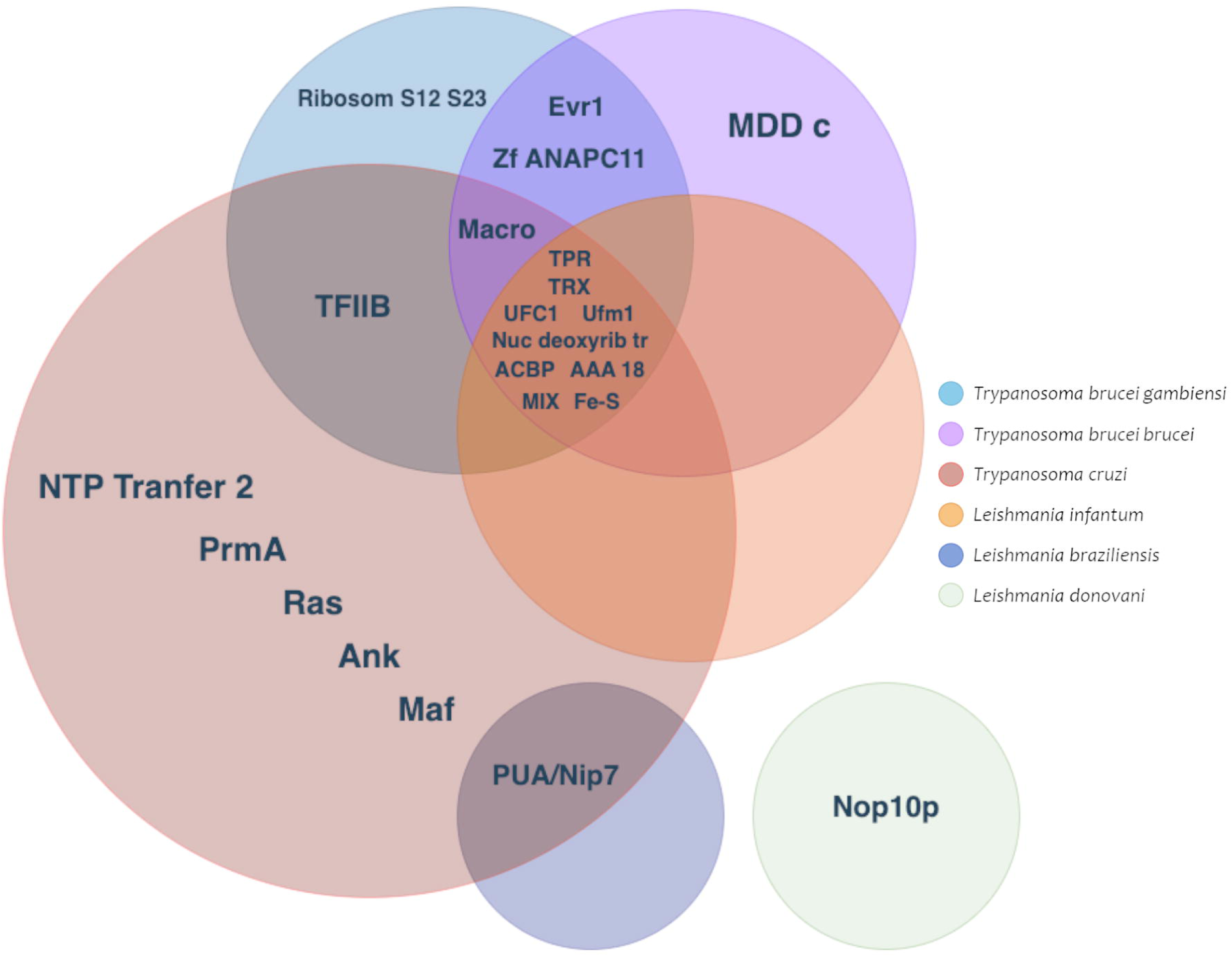
Venn diagram illustrating the shared functional domains among trypanosomatid species. Each circle in the diagram represents a distinct trypanosomatid species, and the predicted functional domains are denoted by bold abbreviations. Our analysis primarily concentrates on the centrally located shared domains, which have the highest number of species located in the middle of the Venn plot.

## Multi-target functional domains identified

### Ubiquitination: UFC1 and Ufm1

In our research, we employed a combination of SEBFP and STBFP methodologies to uncover various functions. However, as we delved into the specifics of each function, it became evident that there were significant gaps in our understanding. For instance, we identified the UFC1 and Ufm1 domains, both of which play roles in the ubiquitination process—a post-translational protein modification system utilizing multiple ubiquitin molecules to regulate cellular functions [61].

The UFC1 domain is a member of the E2-type proteins (conjugators) in the ubiquitination cascade. It plays a vital role in receiving a specific type of ubiquitin (Ub), known as Ubiquitin fold modifier 1 (Ufm1), from Uba5 (E1-type activators) through a thiolester bond mediated by a cysteine [62]. This process is crucial for central nervous system (CNS) development in humans, and bi-allelic mutations in UFC1 have been linked to compromised neurological phenotypes, leading to severe encephalopathy and progressive microencephaly cases [63]. Mutations like Arg23Gln can affect binding to E1 or E3 proteins (ligands), while Thr106Ile impairs the UFC1-Ufm1 complex formation [63].

In trypanosomatids, specifically *T. brucei*, research has shown that the glycoproteins on their surface rely on the ubiquitination process, particularly Invariant Surface Glycoproteins (ISG). After undergoing ubiquitination, these proteins are internalized and degraded, promoting cell surface renewal. This implies that ubiquitination serves as a general mechanism for regulating surface proteins with transmembrane domains [64]. However, research also reveals variations in the machinery involved in this process, such as the absence of orthologs for E3 proteins in the *T. brucei* genome, suggesting alternative mechanisms or factors may be in play in these eukaryotes [65].

Regarding the Ufm1 domain, a study conducted by Yoo et al. in 2014 found that certain ubiquitin ligases are overexpressed in tumors, promoting cell proliferation. This study also identified interactions between estrogen receptors (ERα) and Activating signal cointegrator 1 (ASC1) molecules through the Ufm1 domain, leading to the activation of ERα target genes associated with excessive cell proliferation and breast cancer [66]. The significance of ubiquitylation in the human body underscores the need for further research into UFC1 and Ufm1, especially in the development of targeted inhibitors for specific members of the E3 family [67]. Meanwhile, the proteasome pathway, which is essential for the degradation of cytoskeletal proteins in *T. cruzi*, shows promise as a drug development target. Studies suggest that inhibiting the proteasome can disrupt the evolutionary cycle of *T. cruzi*, making it vital to explore this pathway for new drug targets [68–70]. Hence, it is crucial to persist in the investigation of this pathway to unearth novel drug targets.

### Electron transfer: Fe-S cluster

The Fe-S domain assumes a pivotal role in modulating transcriptional and translational processes within genes [71]. Originating during the prebiotic era on Earth, this domain finds its basis in the abundance of iron and sulfur, coupled with their structural versatility and chemical reactivity, facilitating the formation of Fe-S clusters (iron and sulfur). These clusters function as “molecular switches,” capable of influencing gene regulation at multiple levels [72]. Their intrinsic chemical properties make them active participants in various biological processes, encompassing photosynthesis, DNA replication and repair, respiration, nitrogen fixation, and more. Consequently, proteins containing Fe-S clusters are ubiquitous across all organisms [73,74].

In humans, the cytosolic Fe-S cluster assembly (Cia) targeting complex is constituted by Cia1, MMS19, Cia2A, and Cia2B (also known as Fam96b) [75]. Notably, Cia2A and Cia2B hold significance in iron homeostasis and participate in the maturation of the iron regulatory protein (IRP1) [76]. Fam96a, also recognized as Cia2A in mammals, has demonstrated its role in reducing gastrointestinal stromal tumors, where its interaction with apoptotic peptidase activation factor 1 heightened tumor sensitivity to apoptosis processes [77,78]. Additionally, Fam96a depletion has been linked to organ damage protection and increased survival rates for sepsis patients by reducing the secretion of pro-inflammatory cytokines by macrophages [79].

In trypanosomatids, like *T. brucei*, the presence of the Cia complex has been acknowledged; however, the precise components for Cia in *T. brucei* remain uncertain, and it’s been suggested that these components may have influenced the divergence of the Kinetoplastea class from other eukaryotes [80]. The ISC machinery, responsible for the maturation of cytosolic and nuclear Fe-S proteins, has been proposed to directly impact the formation of flagella in these parasites, particularly under conditions affecting glycolysis and mitochondrial function [81,82]. *Leishmania* spp. also possess Cia components with variations in their mechanisms. Given the conservation and essential role of the Cia pathway in Fe-S cluster assembly within trypanosomatids, it presents a promising source of potential pharmaceutical targets, with Fam96a being one of the identified candidates in *T. brucei* [80] and detected via STBFP analysis (S2 Table).

### Lipid metabolism: ACBP

ACBPs, in general, function by binding to acyl-CoAs, thereby preventing their hydrolysis by hydrolases [83]. This action facilitates the efficient transport of acyl-CoAs to cellular acylation machinery. In humans, ACBPs, also known as ACBDs, have been linked to various cellular processes, including lipid metabolism and cell signaling [84].

In trypanosomatids, ACBPs are believed to protect against inhibition by acyl-CoA esters of acetyl-CoA carboxylase and long-chain mitochondrial adenine nucleotide translocase [85]. These proteins also play a role in the transport and donation of acyl-CoAs for β-oxidation in mitochondria and the synthesis of glycerolipids in microsomes [86].

Experimental studies on trypanosomatids, particularly *T. brucei*, have revealed the significant role of ACBP in the biosynthesis of glycosylphosphatidylinositol (GPI). Specifically, ACBP donates myristoyl-CoA for fatty acid remodeling reactions, causing a conformational change in fatty acids [87]. This alteration allows them to exit the endoplasmic reticulum and anchor to Variant Surface Glycoproteins (VSG), which is crucial for the formation of the VSGs [88]. Once anchored, VSGs undergo further processing in the Golgi complex, including the addition of galactoses, before being transported to the plasma membrane [85].

Furthermore, trypanosomatids rely on GPI anchors for various essential functions, such as evading the host complement system in *T. brucei*, facilitating host cell interaction and sialic acid uptake in *T. cruzi*, and protecting against extracellular hydrolases while interacting with the insect vector’s midgut in *Leishmani*a spp. These membrane components are pivotal for various critical processes, highlighting the importance of understanding the functionality of domains like ACBP in these parasites [89].

### Energy maintenance: AAA18

AAA 18 domain, also known as adenylate kinase (AK), plays a role in regulating cellular energy management by maintaining adenine nucleotide homeostasis. AAA 18 catalyzes the reversible conversion of ATP and AMP molecules into two ADP molecules, contributing to the balance of adenine nucleotides and cellular energy charge, thereby supporting optimal cellular functions and metabolism [84].

Studies have revealed the presence of multiple isoforms of AAA 18 with diverse subcellular localizations and functions. In *T. cruzi*, a cytosolic isoform of AAA 18 has been identified, and it plays a crucial role in the energy metabolism of the parasite, particularly during the growth of the epimastigote form. This suggests a potential correlation between the activity of AAA 18 and the energy demands of the parasite. Interestingly, the expression of the cytosolic AAA 18 isoform decreases when another phosphotransferase, arginine kinase, is overexpressed, indicating a potential functional connection between these two enzymes and a compensatory mechanism for maintaining the parasite’s energy homeostasis [90].

Furthermore, the number of AAA 18 isoforms varies among different species. In the case of *T. brucei*, Ginger et al. (2005) conducted a study that revealed seven isoforms of AAA 18 in this organism. The existence of multiple isoforms suggests a form of metabolic adaptation, leading to increased complexity in energy homeostasis reactions. These isoforms are expressed in both procyclic trypanosomes found in the midgut of the tsetse fly and metacyclic trypanosomes located in the bloodstream, making them particularly relevant in the context of parasite infections [90]. Additionally, in mammals, several AAA 18 isoforms have been associated with cancer, such as the overexpression of AK2 in lung cancer patients, which is linked to the reprogramming of energy metabolism—an essential aspect of cancer progression [91].

Despite our understanding of the functions of ACBP and AAA 18 domains, many mechanisms related to these domains remain enigmatic. One intriguing area of research is the relationship between ACBPs and peroxisomes, given the critical role of peroxisomes in various neurological disorders like Alzheimer’s, autism, and amyotrophic lateral sclerosis [92]. With these proteins playing a significant role in humans, concerns arise when targeting them with drugs due to potential systemic consequences. Therefore, further research is essential to enhance our understanding of the selectivity and implications of targeting ACBPs and AAA 18 in the context of human health.

### Purine metabolism: TRX

TRX domain is a pivotal component of the purine metabolism pathway. Purines play a crucial role in the synthesis of signaling molecules like cyclic adenosine 3,5-monophosphate (cAMP) and are integral to the cellular energy system, including adenosine triphosphate (ATP). Furthermore, they participate in the production of RNA and DNA, working in tandem with pyrimidines [93,94]. However, experiments conducted on *T. brucei* cultures have raised questions about the indispensability of this protein due to the presence of a second TRX enzyme, TRX2 [95].

Significant differences in conservation between TRX2 and TRX1 exist among various trypanosomatid species. TRX1 exhibits a lower degree of conservation, with approximately 32% identity between *T. cruzi* and *T. brucei* [95]. However, our alignment results indicate a higher identity of 50% between these species, with the lowest being 44% between Tcr and Lif. This supports the hypothesis that the domain we’ve identified is TRX1. Studies have revealed that the absence of TRX2 leads to compromised parasite proliferation in *T. brucei*, while the deficiency of TRX1 doesn’t impact parasite proliferation [95]. Consequently, it appears that TRX1 may not be a promising target for drug development.

In the upcoming sections, we will explore several protein domains that we have identified as promising candidates for multi-target studies in the development of novel drugs. These domains play critical roles in Trypanosomatids, making them valuable subjects for therapeutic investigation. To gain a comprehensive understanding of these domains, we conducted extensive analyses, including the generation of their oligomeric structures using AlphaFold2, rigorous evaluation of quality parameters, and the prediction of druggable sites. Detailed sequences of these domains are available in the S3 table, which will aid in further elucidating their potential as therapeutic targets.

### Cyclogenesis: IFT – TPR

The tetra-trico peptide repeat domain (TPR) is a highly conserved structure comprised of multiple repeats of 34 hydrophobic amino acids, found in various organisms [96]. Its involvement in critical processes, such as transcriptional inhibition, stress response, and mitochondrial protein transport, positions it as a promising target for drug development [97–99].

Our evaluation of the TPR’s conservation across trypanosomatid species revealed high levels of identity, ranging from 73% (between Lif and Tbg/Tbr) to 98% (between Tbg and Tcr). Furthermore, the human isoforms of IFT70 (A and B) showed an identity range of 51% to 54% (Fig 4A). While the isoforms share a considerable degree of similarity, it’s essential to emphasize the significance of the IFT train’s involvement in various processes, which we will describe in detail below.

**Fig 4.**
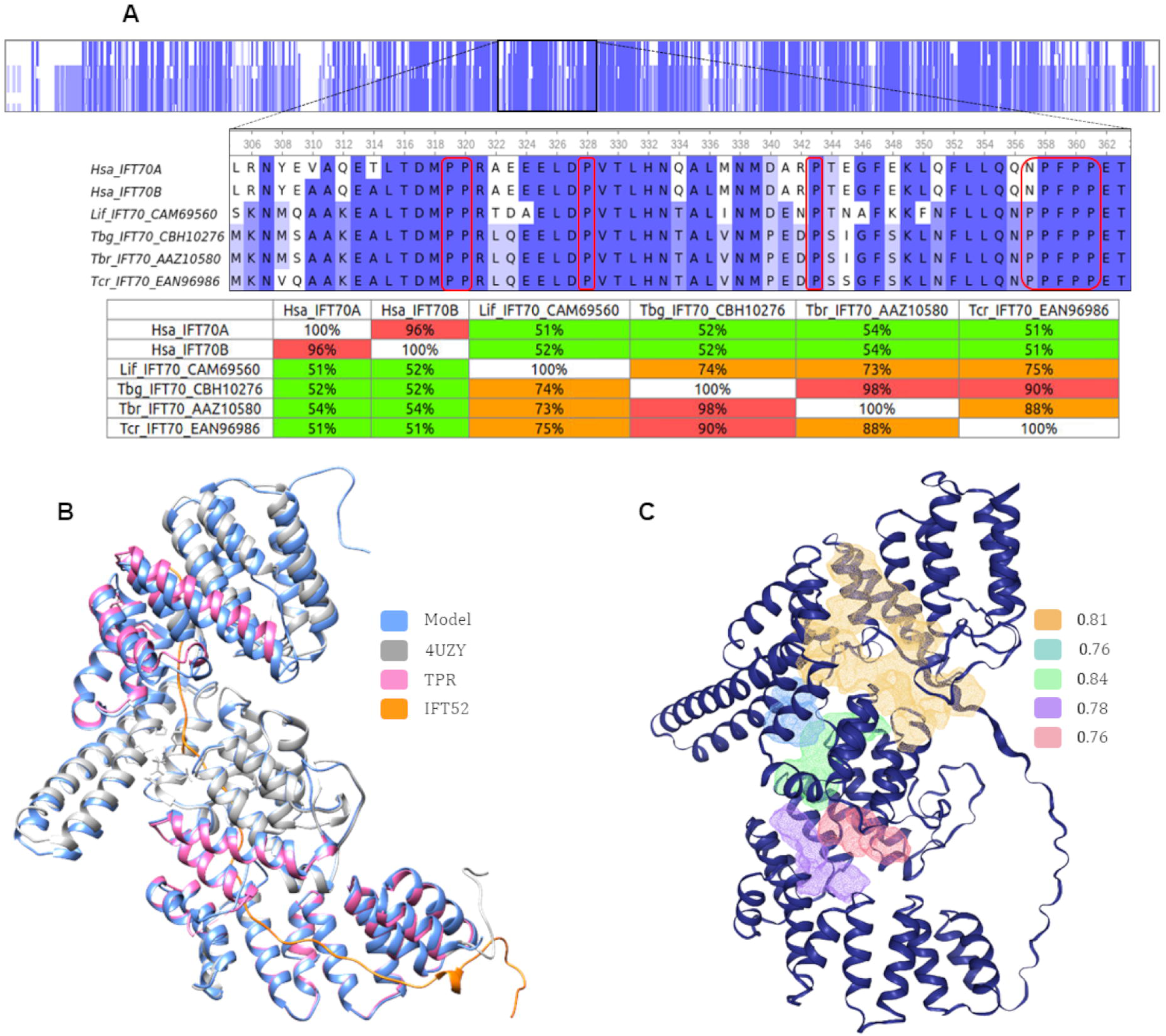
TPR – Sequential and structural analysis. (A) Shows the alignment of trypanosomatid and human (Hsa) sequences, accompanied by a graphical summary revealing the percentage of identity. Notably, the proline-rich region is highlighted, demonstrating its conservation across species. The alignment matrix below employs color-coding within the UGENE software, indicating dissimilarity percentages: white (0%), red (10%), orange (25%), green (50%), blue (70%), and gray (90%). (B) Illustrates the alignment between the 3D model generated by MHOLline for the Lif sequence (dark blue) and the 4UZY template (gray). The TPR domain is depicted in pink, while the IFT52 protein, resolved with the PDBid: 4UZY structure, is represented in orange. (C) Results for the druggability prediction where in a blue cartoon is IFT70 of *L. infantum* (Lif) and the druggability score and corresponding surface area values are color-coded: green (0.84, 888.13 Å²), orange (0.81, 3311.56 Å²), purple (0.78, 878.57 Å²), pink (0.76, 746.52 Å²), and blue (0.76, 605.84 Å²).

The TPR domain’s versatility in cellular location, being found in various compartments such as the nucleus, cytoplasm, and mitochondria [96], was confirmed in our Psort prediction for trypanosomatid species. Tcr, Tbr, and Lif were predicted to be mitochondrial, while Tbg was cytoplasmic. The TPR domain’s role in mediating protein-protein interactions and assembly of protein complexes was underscored by our Structure-Based Functional Prediction (STBFP) results, which identified the IFT70 protein (PDBid: 4UZY) [100]. IFT70 is present in the IFT train which is a crucial component of the intraflagellar transport protein complex responsible for cyclogenesis, an evolutionarily conserved transport process involving the bidirectional movement of particles within cilia [101].

As noted in the structural analyses, the 4UZY template corresponds to the IFT70 protein complexed with part of IFT52 from the organism *Chlamydomonas reinhardtii* (PDBid: 4UZY). The interaction between IFT70 and the IFT52 protein occurs through a proline-rich region [102], and we observed that most of these regions are conserved, even in humans (Fig 4A). Therefore, the tertiary and primary structures exhibited significant similarity to a functionally characterized protein (Fig 4 A and B), reinforcing the connection between structure and functionality in proteins.

To enhance the accuracy of our structure prediction, we employed AlphaFold2 to generate new models for each species, taking into account the oligomeric states of the proteins. The results for IFT70 can be found in S1 Fig and to ensure the quality of these models, conducted evaluations of the models using MolProbity [56] (Clash score, MolProbity score, and Ramachandran favored regions), as well as ProSa [57] (as shown in S4 Table). Notably, the best IFT70 model was identified for the *L. infantum* (Lif) organism (accessible via this ZENODO link: https://doi.org/10.5281/zenodo.10072065). Consequently, we employed this model to predict potential binding sites using DoGSiteScorer [58], which assigns druggability scores within the range of 0 to 1. Higher scores indicate more favorable druggable pockets, guiding our search for potential drug-binding sites. The results for the Lif model indicate the presence of five promising regions (Fig 4 C), with the majority of them located within the IFT52 binding site, reaffirming the crucial role of this region in the functionality of IFT70.

Furthermore, the significance of the IFT52 protein in stabilizing the IFT-B1 complex and its direct involvement in cilia formation has been well-documented in studies across various organisms [103,104]. For instance, the study by HUET et al. in 2019 demonstrated that other proteins within the IFT train, such as IFT172 (IFT-B2), were pivotal for proper cyclogenesis and flagellar organization in *T. brucei*. These findings underscore the indispensable role of the IFT train in cyclogenesis and emphasize the critical contributions of individual proteins, like IFT52, in maintaining the complex’s stability and functionality.

Despite the significant progress in understanding the IFT train, questions remain regarding the interactions and subunits involved in complex stabilization. However, the potential of the IFT70 protein and other IFT train proteins as targets for drug development in trypanosomatids is promising, given their pivotal role in critical cellular processes and preliminary in vitro study results. Further investigations may reveal new drug development opportunities.

### Nucleosides transferases: Nuc deoxyrib tr

Nucleoside 2’-deoxyribosyltransferases (NDTs), also known as Nuc deoxyrib tr according to domain databases’ nomenclature, constitute a protein family that mediates the transfer of the 2’-deoxyribosyl group between 2’-deoxyribonucleosides (donor) and nucleobases (acceptor) [105]. These enzymes belong to the class of Nucleoside Phosphorylases (NPs), which are further divided into ribo– and 2’-deoxyribonucleosides. The class encompasses various subclasses, including type I NDTs (PDT), responsible for deoxyribotransfer between purines only, and type II NDTs (NDT), which enable transfer between purines and/or pyrimidines [106]. NDTs have been identified in various species, including parasitic ones like *T. brucei*, *L. major*, and *L. mexicana*, as well as in lactobacillus species like *Lactobacillus helveticus* [107,108].

Bosch et al., 2006 conducted a comparative analysis of NDTs among different trypanosomatid species, including *T. cruzi*, *L. major*, *L. infantum*, and *T. brucei*, along with a sequence from Lactobacillus helveticus. The study revealed that the percentage of identity between *T. brucei* and other species ranged from 84.2% (*T. cruzi*) to 89.5% (*L. major* and *L. infantum*), and only 21% with *L. helveticus*. Our analysis of predicted NDT domain sequences showed identity ranging from 65% between Tcr and Lif to 100% between Tbg and Tbr (Fig 5A).

**Fig 5.**
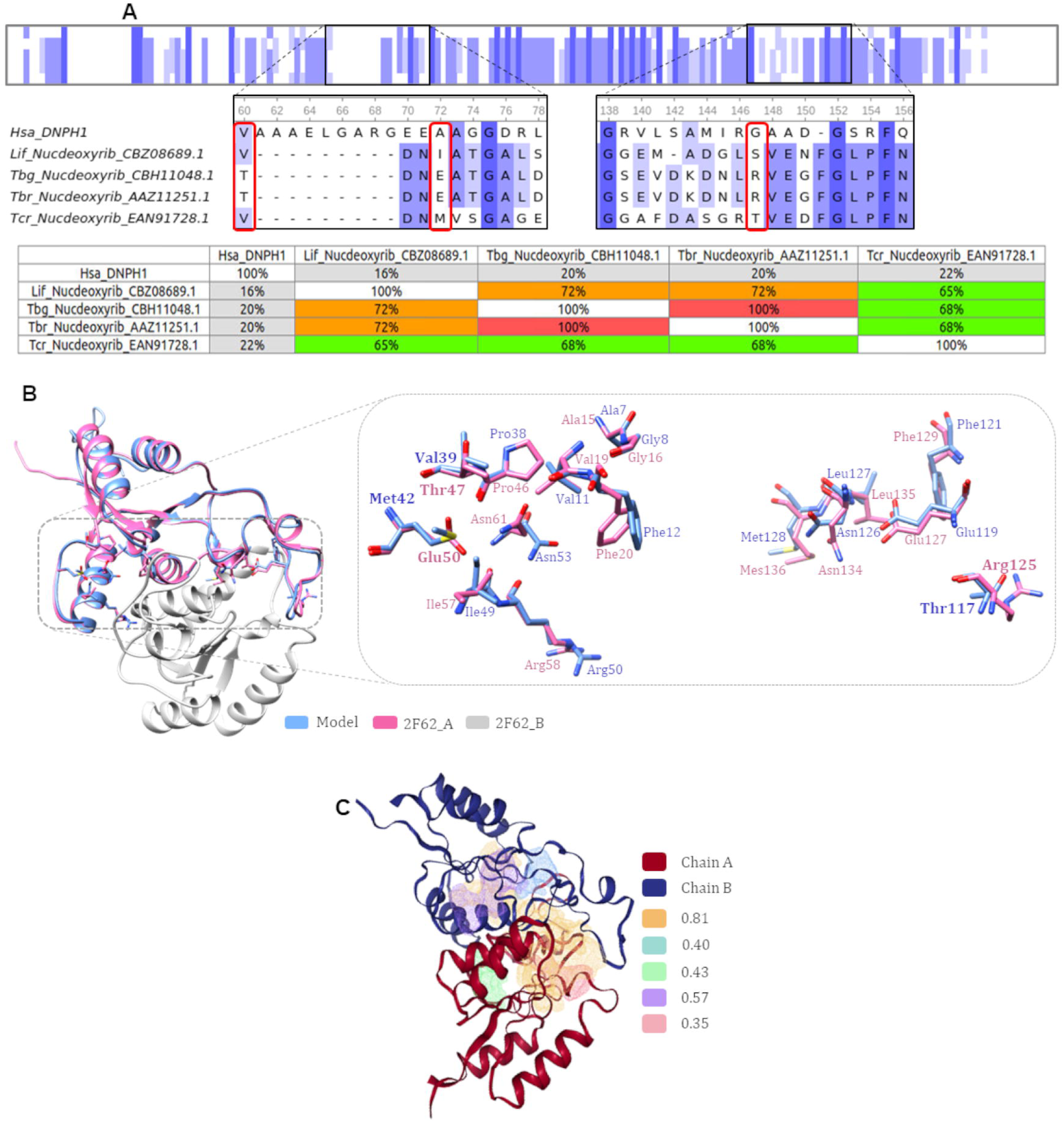
NDT – Sequential and structural analysis. (A) The alignment reveals the comparison between trypanosomatid and human sequences. Notably, the highlighted region contains three amino acid mutations. The right side of the figure displays the aligned residues: Ala15, Gly16, Val19, Phe20, Pro46, Thr47, Glu50, Ile57, Arg58, Asn61, Arg125, Glu127, Phe129, Asn134, Leu135, and Met136. Among these residues, three mutations were observed: Thr47Val, Glu50Met, and Arg125Thr, as indicated in bold. The numbering of *T. cruzi* amino acids was adjusted to account for the eight fewer amino acids compared to the template. (B) The alignment between the predicted model for the Lif protein (light blue) and the template with PDBid: 2F62 (chain A in pink and chain B in gray). This template was selected due to having more mutations and the lowest RMSD. The aligned residues match those presented in part (A) and highlight the conserved regions among the sequences. (C) Results for the druggability prediction where Nuc deoxyrib tr Lif is in red (chain A) and blue (chain B) in a blue and the druggability score and corresponding surface area values are color-coded: orange 0.81 (1649.44 Å²); purple 0.57 (354.17 Å²); green 0.43 (231.66 Å²); blue 0.4 (264.39 Å²); pink 0.35 (213.49).

Bosch’s work also highlighted conserved regions among trypanosomatids, which included key residues within the 2’-deoxyribose binding site, such as Ala15, Gly16, Val19, Phe20, Pro46, Thr47, Glu50, Ile57, Arg58, Asn61, Arg125, Glu127, Phe129, Asn134, Leu135, and Met136 [108]. Comparing these regions among the generated structures, it was evident that most of them were conserved across species, with only a few mutations observed, such as Thr47Val in Tcr and Lif; Glu50Ile and Arg125Ser in Lif; Glu50Met and Arg125Thr in Tcr (Fig 5B). The RMSD values between the mold and the models ranged from 0.467 Å (Tcr) to 0.611 Å (Lif).

Following the utilization of AlphaFold2 and the subsequent evaluation of model quality (refer to S1 Fig and S4 table for details), we proceeded to select the Lif structure for predicting druggable sites. Among the identified sites, the orange region (Fig 5C) scored the highest, signifying its potential druggability. Notably, these druggable sites are concentrated within a region that contains key amino acids critical for the proper functioning of this protein.

Analysis in parasites such as *Trypanosoma brucei*, *Trypanosoma cruzi*, and *Leishmania major*, showed that Nucleoside 2’-deoxyribosyltransferases (NDTs) play a crucial role in the processes of recycling and recovery of purines, as well as in the de novo biosynthesis of purines [108]. These parasites employ various enzymes to remove ribose or deoxyribose from purine nucleosides, making it challenging to determine which enzymes are essential in the purine turnover pathway.

In contrast, humans rely on Deoxynucleoside 5-monophosphate N-glycosidase (DNPH1 – EC 3.2.2.) for similar functions, but differences in substrate specificity and essentiality differentiate it from NDTs [109]. Furthermore, it has been suggested that DNPH1 is not an essential gene, as many organisms can survive without it. These findings support the idea that NDT and DNPH1 enzymes are distinct. To further compare the two, we aligned the predicted NDT domain sequences of various species with the DNPH1 sequence of Homo sapiens (UniProtKB code: O43598). The identities between these sequences ranged from 16% (between Lif and Hsa) to 22% (between Tcr and Hsa), confirming their significant differences (Fig 5A).

Given the precariousness of enzymes involved in de novo purine biosynthesis in *T. brucei* and their reliance on the host for purine uptake, inhibiting NDT, an enzyme crucial for their processes, could significantly impact trypanosome replication [108]. In light of the similarity among trypanosomatid sequences and the absence of NDT in humans, NDT presents a promising therapeutic target for further exploration.

### Mitochondrial membrane: MIX

The MIX domain is a unique protein structure anchored to the mitochondrial membrane and found exclusively in kinetoplastids [110], a group of organisms that includes Trypanosoma and Leishmania species. In these kinetoplastids, the precise mechanisms of mitochondrial protein import and processing are not fully understood. One approach to gaining insights into the function of the MIX domain is to study its association with the Cytochrome C Oxidase (COX) multiprotein complex, as proposed by Gorman et al. in 2011 [111].

Cytochrome C Oxidase (COX) plays a crucial role in mitochondrial cellular respiration, driving oxidative phosphorylation to generate ATP molecules. This energy production process involves various protein complexes (I, II, III, IV, and V) [112], with COX representing complex IV in trypanosomatids [111]. Notably, the COX complex in these organisms exhibits distinctive features in 8 of the 10 smaller subunits, for which no counterparts are identified outside the Kinetoplastida group [113]. Early hypotheses suggested that the MIX domain may be instrumental in directing or assembling COX components [111].

The MIX domain comprises small sequences characterized by a conserved primary structure, facilitating their targeting of mitochondria in parasitic and yeast cells [110]. In their 2011 study, Gorman et al. compared the MIX sequence from *Trypanosoma brucei* with that of other trypanosomatids, specifically *Leishmania major* and *Trypanosoma cruzi*, revealing identity percentages ranging from 75% to 66%, respectively. In the present study, comparisons of sequences predicted to contain this domain show identity percentages ranging from 67% to 99% (Fig 6A).

**Fig 6.**
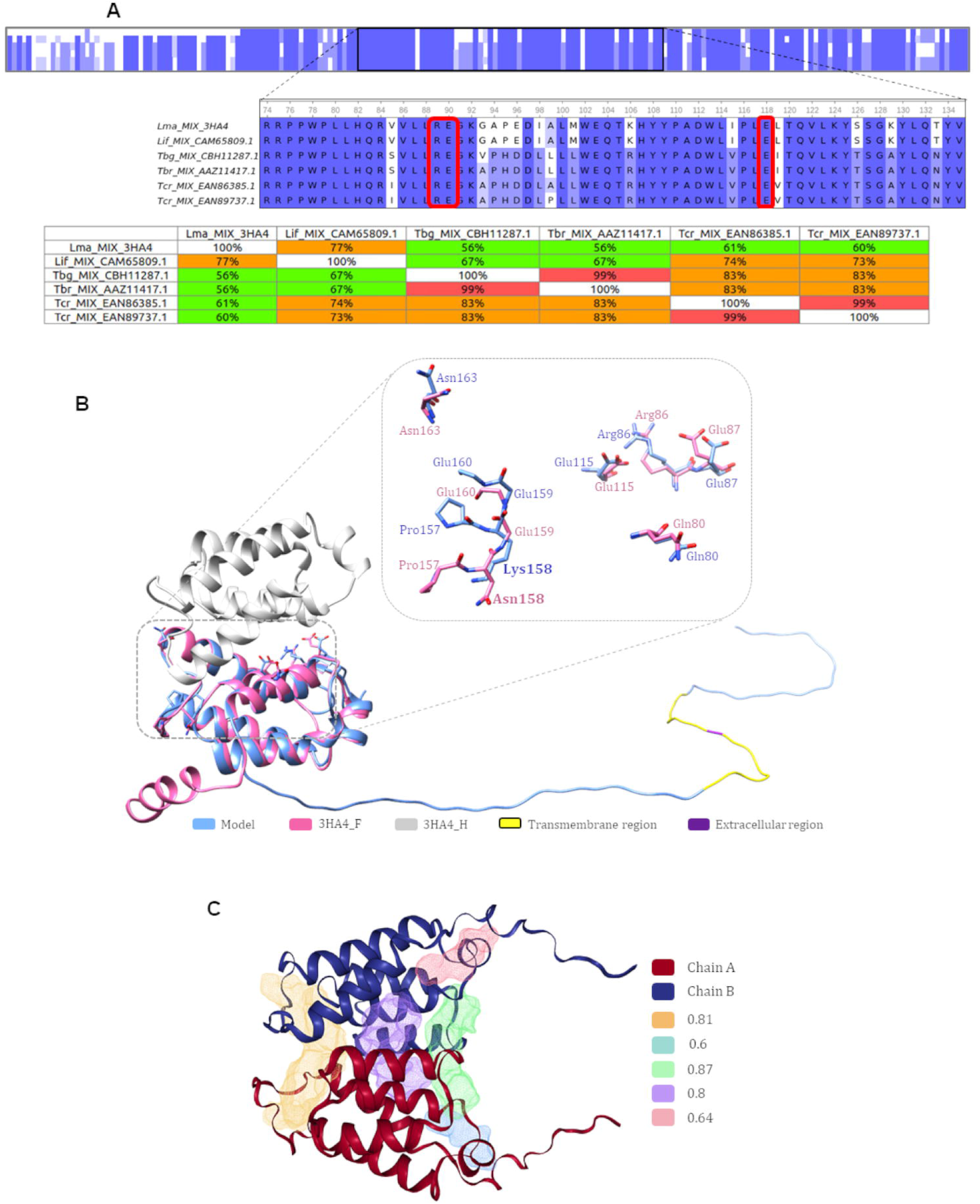
MIX – Sequential and structural analysis. (A) This section features the alignment of trypanosomatid sequences, with a specific emphasis on critical amino acid residues (Arg86, Glu87, and Glu115) that play a pivotal role in protein dimerization. (B) The structural alignment showcases the model sequence CAM65809 from *L. infantum* in light blue, contrasted with the template PDBid: 3HA4 (F chain in pink and H chain in gray). Notably, this template contains the sole mutation found in Asn158Lys, represented in bold. Additionally, predictions from the DeepTMHMM web server are illustrated, highlighting the transmembrane region in vibrant yellow and the extracellular region in regal purple. A display of amino acid residues is provided on the right side of the image, featuring elements like salt bridge formers (Arg86, Glu87, and Glu115), as well as residues that engage in hydrogen bonds without mediation by water molecules (Gln80, Glu160, Pro157, Gly159, and Asn163). (C) The results of druggability assessments for Mix Lif are presented in red (chain A) and blue (chain B), utilizing a cartoon representation (excluding the initial 70 amino acids). Druggability scores and their corresponding surface area values are color-coded: green 0.87 (602.82 Å²); orange 0.81 (1446.28 Å²); purple 0.8 (772.76 Å²); pink 0.64 (404.49 Å²); blue 0.6 (369.83 Å²).

Moreover, Gorman et al. in 2011 identified three critical regions for the MIX domain’s structural integrity in various trypanosomatid species, using *Leishmania major* (Lma) as the model organism. The first region involves three amino acid residues (Arg86, Glu87, and Glu115) essential for dimerization. The second region consists of amino acid residues that form salt bridges (Arg89, Glu90, and Glu118), while the third region comprises amino acid residues that engage in hydrogen bonding without water molecule mediation, including Gln80, Glu160, Pro157, Asn158, Gly159, and Asn163.

In our own work, we compared the same Lma sequence used by Gorman et al. in 2011 and found identity percentages ranging from 56% (between Lma and Tbg/Tbr) to 77% (between Lma and Lif). The first two regions remained conserved, while we identified a single mutation in the third region, specifically in Lif: Asn158Lys (Fig 6B). This mutation may impact the formation of chemical bonds due to the distinct physicochemical characteristics of the lysine radical, suggesting the need for further investigation to understand its effects on hydrogen bond formation.

In our structural analyses, the STBFP results and the template used led to the structure PDBid:3HA4, representing a MIX protein from the *L. major* organism [114]. However, we observed that the generated models exhibited a significant region without a clearly defined secondary structure, encompassing residues 1 to 70. To address this issue, we employed the transmembrane region predictor, DeepTMHMM, based on prior descriptions of the MIX domain. The results indicated a consensus for the Lif and EAN89737 species of Tcr, while transmembrane regions were not predicted for Tbg and Tbr (Table 2). This provides valuable insights into the structural characteristics of the MIX domain in different trypanosomatid species.

**Table 2.** Results DeepTMHMM.

| Organism | Regions | Prediction |
| --- | --- | --- |
| **Lif** | 1 – 22 | I |
|  | 23 – 32 | Tmhelix |
|  | 33 | O |
|  | 34 – 43 | Tmhelix |
|  | 44 – T5 | I |
| **Tbg** | 1 – 198 | I |
| **Tbr** | 1 – 198 | I |
| **Tcr** | 1 – 27 | I |
|  | 28 – 37 | Tmhelix |
|  | 38 | O |
|  | 39 – 48 | Tmhelix |
|  | 49 – 198 | I |

<sup>a</sup>i: Intracellular region.
<sup>b</sup>TMhelix: Transmembrane region.
<sup>c</sup>o: Extracellular region.

Uboldi et al. (2006) postulated the existence of an N-terminal amphipathic α– helix within the MIX domain, followed by a subsequent transmembrane region, with the transmembrane portion typically being laterally inserted into the mitochondrial membrane. In the case of *L. major*, this transmembrane region consists of the initial 45 residues [110]. The DeepTMHMM results for proteins that possess the MIX domain, even those with unknown functions, align with this characteristic.

In terms of the structural attributes of MIX, it comprises seven α-helices that collectively form the coiled domain [114]. This pattern of folding is consistent with our generated models, and the root mean square deviation (RMSD) values derived from the comparison of these models with the template (PDBid: 3HA4) ranged from 2.069 Å (Tbr) to 2.493 Å (Lif). It is worth noting that the variability in RMSD values is likely associated with the N-terminal segment (residues 1-45), for which the template lacks a well-defined structure (Fig 6B).

In summary, the MIX domain is universally expressed across all evolutionary stages of Leishmania, including the amastigote form residing in the mammalian host. Intriguingly, both in vitro and in vivo experiments have demonstrated that the deletion of two MIX gene alleles leads to a reduction in infectivity and virulence [110]. While initial assessments in *T. brucei* suggested that COX activity was solely required in the epimastigote form, indicating that the interaction between MIX and COX was specific to this developmental stage [110,114], subsequent research has unveiled the presence of MIX in the blood stage. This suggests that MIX could engage with other proteins and catalyze additional reactions [111]. Given the unique characteristics of MIX within the Kinetoplastida group and its pivotal roles in diverse processes, it emerges as a promising molecular target for the development of novel drugs.

## Conclusion

Based on the SEquence-Based Functional Prediction (SEBFP) and Structure-Based Functional Prediction (STBFP) results, we were able to predict functional domains for 67 trypanosomatid proteins, of which nine domains were shared among the majority of trypanosomatid species. These domains were analyzed, by consulting bibliographic resources, and we were able to clarify their functions and their existing correlations with trypanosomatids.

Among the domains analyzed, we found those that had several isoforms, not being crucial for the parasites under study, TRX, ACBP, and AAA 18 [95,115][115]. Additionally, the complexity of the pathways in which they are inserted, along with the lack of detailed information on all the inherent metabolic processes, currently hinders the development of inhibitors for these targets.

In addition to the aforementioned domains, we also identified two domains that are integral to the ubiquitination process, a crucial mechanism for protein degradation. Research has shown that mutations in UFC1, for instance, can result in neurological impairment [63]. Similarly, Ufm1 can recruit transcriptional coactivators in humans, contributing to excessive cell proliferation and ultimately leading to conditions such as breast cancer [66]. Both domains possess complex intricacies that need to be addressed before considering them as therapeutic targets. Nonetheless, the proteasome, another component of the ubiquitination pathway, has been studied extensively due to its significant role in *T. cruzi* [116]. Inhibitors of the proteasome have already been tested, and the results demonstrate impairment of the infective evolutionary form, indicating that a study considering the proteasome as a therapeutic target could yield promising results.

Lastly, we identified domains that can be essential for the survival and virulence of the trypanosomatids under study, namely Nuc deoxyrib tr, MIX, and TPR. Nuc deoxyrib tr is critical in de novo purine biosynthesis, and studies on *T. brucei* have shown that these parasites do not produce an abundance of such proteins, making them heavily reliant on the catalytic activity of this domain for survival [108]. Regarding the MIX domain, this is exclusive to kinetoplastids and *in vitro* and *in vivo* tests demonstrate that its deletion led to a reduction in infectivity and virulence [111]. Furthermore, we also discovered the TPR domain, specifically the IFT70 protein, which is involved in a protein complex that facilitates intraflagellar transport. IFT70 is a vital member of the IFT train, and studies on *T. brucei* have confirmed that the train is necessary for the transportation of ciliary assembly precursors from the cytoplasm into the cell body [103]. Therefore, we believe these targets hold promise for multi-target studies aimed at developing innovative therapeutic approaches.

## Supporting information

S1 Table Domain index

S2 Table Functional prediction results

S3 Table Proteins Domains sequences

S4 Table Oligomeric AlphaFold Models Validation

## Acknowledgments

RSLR expresses sincere gratitude to the Graduate Program in Computational and Systems Biology at Instituto Oswaldo Cruz for their unwavering support. Furthermore, she would like to extend her heartfelt appreciation to Instituto Oswaldo Cruz and the Capes program for generously providing financial assistance (No. 88887.596358/2020-00, No. 88887.801809/2023-00).

## Supporting Files

**S1 Fig.** AlphaFold2 prediction. Presented in both cartoon and surface views. These models pertain to the TPR, Nuc deoxyrib tr, and MIX domains, covering various species, namely *L. infantum* (Lif), *T. brucei gambiensi* (Tbg), *T. brucei brucei* (Tbr), and *T. cruzi* (Tcr). In this representation, monomeric TPR models are depicted in vibrant purple, offering insights into the individual subunits. Dimeric structures are presented with chain A in a distinct orange shade and chain B in a contrasting blue hue, allowing for clear differentiation between the two subunits. Arrows within the figures highlight regions where MIX transmembrane segments are predicted, indicating potential areas of interest for further study and analysis.

**S1 Table. Domain index**.

**S2 Table. Functional prediction results.** The abbreviations in the table mean WR – Without Results; UP – Uncharacterized Protein; UCP – Uncharacterized Conserved Protein.

**S3 Table. Proteins/Domains sequences.**

**S4 Table. Oligomeric AlphaFold Models – Validation.**

